# An fMRI study of composition in noun and verb phrases

**DOI:** 10.64898/2025.12.17.694853

**Authors:** Laurent Bonnasse-Gahot, Doug Bemis, Martin Perez-Guevara, Stanislas Dehaene, Christophe Pallier

**Affiliations:** Centre d’Analyse et de Mathématique Sociales, CNRS, EHESS, PSL University, Paris, France; Cognitive Neuroimaging Unit, Neurospin, INSERM-CEA, 91191 Gif-sur-Yvette, France; Collège de France, Paris, France; Centre National de la Recherche Scientifique, Paris, France

**Keywords:** fMRI, Language, Composition, Syntax

## Abstract

How do the language areas of the human brain combine multiple words into meaningful phrases and sentences remains ill-understood. Here, to address this question, we determined the response profile of temporal and inferior frontal language areas to the composition of up to four words into phrases. We tested whether brain activity increases with the number of merged words, and whether this profile differs for noun and verb phrases. To this aim, we used fMRI to quantify the brain responses to individual noun and verb phrases of varying length and to tightly matched word lists. Increasing phrase length was associated to an increase in activation in all regions of the temporo-frontal language network. The effect was more pronounced for phrases built around verbs than for phrases built around nouns, suggesting that verbs involve a more complex syntactic tree structure than nouns. Even with word lists, several regions, notably the inferior frontal gyrus (IFG) pars triangularis and opercularis and the posterior superior temporal sulcus showed clear increases in activity with the length of sequences, although the words could not be merged into phrases. By contrast, other regions (IFG pars orbitalis, anterior temporal lobe, temporo-parietal junction) did not react to scrambled word lists. Those different functional response profiles inform theories of how composition is implemented in the human brain.

## Introduction

Many brain imaging experiments have sought to identify the brain circuits involved in merging words into phrases during sentence comprehension (e.g. Mazoyer et al., 1993; Pallier et al., 2011; Goucha & Friederici, 2015; Price et al., 2015; Zaccarella et al., 2017; Matchin et al., 2017). For example, Pallier et al. (2011) reported that neural activity increased with the size of constituents (syntactically and semantically coherent chunks of words) in most of the temporofrontal language network. Moreover, when content words were replaced with meaningless pseudowords (creating so called “Jabberwocky” stimuli), activation increased with constituents’ size solely in the Inferior Frontal Gyrus (IFG) and in the posterior Superior Temporal Sulcus (pSTS), suggesting that these regions are implicated in syntactic processing, in agreement with many previous studies (e.g. Rodd et al., 2010; Santi & Grodzinsky, 2010; Tyler et al., 2008; Pattamadilok et al., 2016).

While Pallier et al. (2011) used phrases of up to 12 words, Bemis & Pylkkänen (2013) presented participants with simpler two-words stimuli that could either merge into a simple phrase or not (e.g. “red boat” vs. “car boat”). They observed a stronger response to phrases than to non-phrases in the anterior temporal lobe (ATL). This suggests that this region plays a role in combining meaningful content words, or in the word of the authors, performed “semantic composition”. Consistent with this interpretation, Pallier et al. (2011) reported in the anterior temporal region an effect of constituent size only for normal words but not for Jabberwocky (but see Humphries et al., 2006, who reported structural effect with Jabberwocky stimuli in the ATL).

The interpretation of such patterns of activation is a matter of debate. A recurring question concerns the relative roles of syntactic and semantic combinatorial processes in the patterns of activation detected when comparing sentences to word lists, using Jabberwocky or not. One difficulty is that the boundary between syntax and semantics is not clearcut. Pseudowords in Jabberwocky stimuli are created to avoid mapping on already existing words and lack reference or definition; nevertheless, they also disrupt syntactic processing because it is not immediately evident which part of speech they belong to. Furthermore, Jabberwocky may not fully eliminate semantic-level composition, as many participants mention that the pseudowords evoke a tentative meaning. Goucha & Friederici (2015) argued that derivational morphemes, which are often preserved when generating Jabberwocky stimuli, carry semantic information, and that a pure syntactic contrasts requires removing all derivational morphology from Jabberwocky pseudowords. When they compared the activations evoked by structured vs. random Jabberwocky *without derivational morphology*, only BA44 was involved, in contrast to a situation in which derivational morpheme were kept and more brain areas were involved (see also Zaccarella & Friederici, 2015). Does this mean that BA44 is a pure syntactic area? We would argue that it is still not obvious that pure Jabberwocky sentences are devoid of semantics as one can clearly compute the logical form of a Jabberwocky sentence. In the end, we reckon that whether the logical form is part of semantics or syntax is essentially a matter of terminology and that it more fruitful to identify which exact operations are performed rather than using too general terms.

Another question concerns the nature of the processes uncovered by comparing structured versus random lists. Stronger activations to well-formed sentences could reflect a variety of processes including syntactic parsing, but also semantic composition, contextual integration, etc. For example, Matchin et al. (2017) interpret their results in terms of top-down syntactic predictions that facilitate sentence processing rather than structure building operations. Note, however, that top-down syntactic expectations must rely on the building up of structures, as, for example, occurs during top-down parsing (e.g. Hale, 2014).

The bulk of the argument of Matchin et al. (2017) lies in the fact that in their experiment, short (2-word long) phrases did not elicit stronger activations than random word lists; only complete sentences (where prediction can come into play) did. Neither does the data from Pallier et al. (2011) show a clear effect of two-word phrases versus lists. On the other hand Bemis & Pylkkänen (2013) report an effect of 2 word phrases, but in the anterior lobe rather than in the IFG.

In order clarify the facts, we decided to focus on short phrases (up to 4 words), contrasting them to matched list conditions where words cannot not be merged into larger chunks. Our aim was to quantify how the activation varies with the number of words, both in phrases and in word lists. Contrary to the design of Pallier et al. (2011) and in line with Bemis and Pylkaanen (2013), the “phrase” stimuli featured a single phrase rather than a series of phrases of constant size, simplifying the prediction of activations patterns. Also, we were interested in separating the case of phrases built around nouns and verbs because different responses to noun and verb phrases have been reported, notably by Tyler et al. (2008). Verbs often have a greater syntactic complexity than nouns, for instance because they take multiple arguments, and linguists postulate that verb phrases involve a greater hierarchy of nodes in the “left periphery”, such as tense and focus nodes, which would predict that verb phrases should induce more intense activations than noun phrases in the brain’s language network.

We measured brain activity generated during the comprehension of progressively larger linguistic structures, ranging from one to four words. In order to control for activity related to non-compositional lexical processing, we also measured activity generated during the comprehension of one to four words, presented in a list context. In past work, the sentence and word list conditions have not always been precisely matched in lexical content. In order to perfectly match these conditions in terms of the specific words they contained, we employed two different phrase types in the phrase conditions, noun phrases and verb phrases, and we constructed the list conditions by interleaving words from different phrase types (see Fig. 1). For example, from the phrases “avec son beau clavier” (*with his nice keyboard*) and “Adam doit plus nager” (*Adam must swim more*), we interleaved the words to generate the associated list conditions “avec doit beau nager” (*with must nice swim*) and “Adam son plus clavier” (*Adam his more keyboard*), where successive words can never combine into a well-formed phrase. Thus, the phrases and lists contained exactly the same words at exactly the same positions. For both trial types, words were presented sequentially and followed by a delay period, after which subjects were asked to judge whether a given probe word had appeared in the trial.

**Figure 1.**
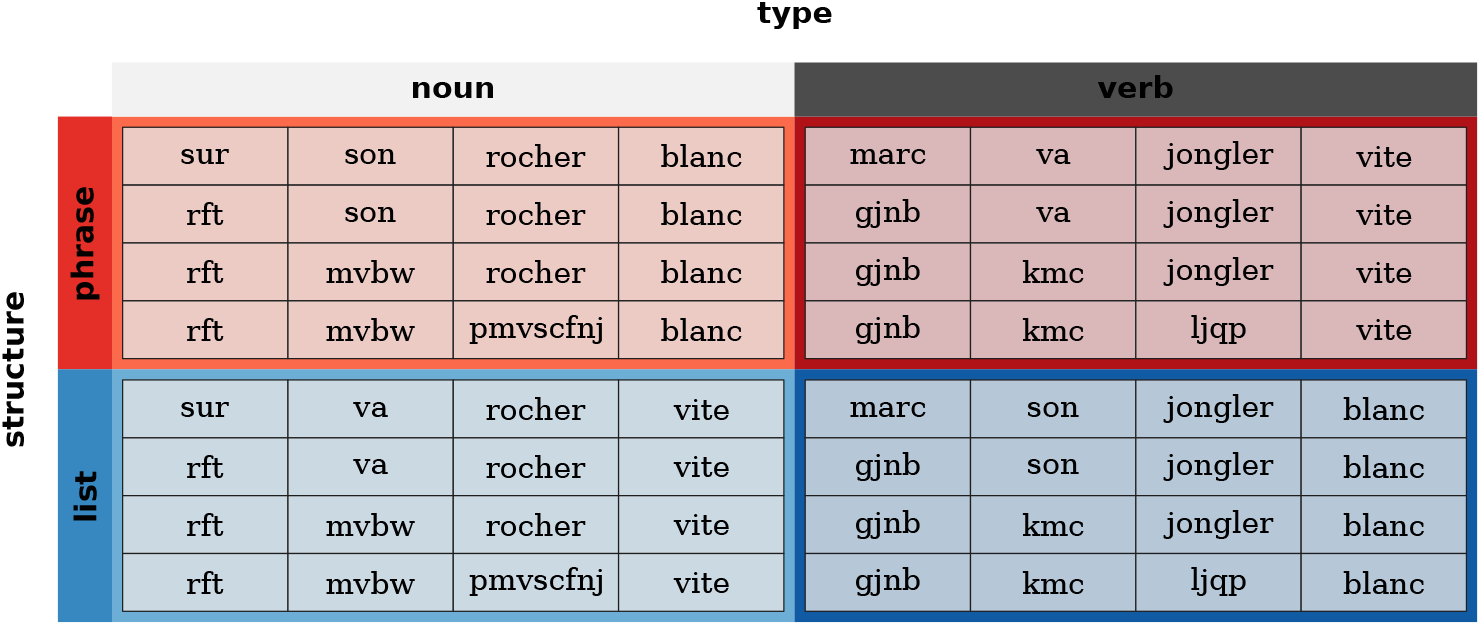
Experimental Design. In each trial, a sequence of consonant strings followed by one to four words was presented. This table shows the 16 conditions generated by a factorial design with three fully crossed factors, hereafter termed Structure (Phrase vs. List), Type (Noun vs. Verb) and Length (1 to 4 words). Note that in the actual stimuli presented during the experiment, an additional consonant string was presented at the start of each sequence.

## Materials and Methods

### Participants

Eighteen right-handed native French speakers, with normal or corrected to normal vision, aged between 20 to 35 years, participated in the experiment. The experiment was approved by the Regional Ethic Committee (Comité de Protection des Personnes du Kremlin-Bicêtre). All participants gave their written consent. Three participants were excluded from the study because of their very poor performance (error rates larger than 20% in the probe detection task where only 1 to 4 words had to be memorized), suggesting that they did not pay enough attention to the stimuli; the average error rate of the 15 remaining subjects was 3.2%.

### Stimuli

Each trial consisted in a series of five successive letter strings. The first stimulus was always a 3 to 8 letters long, non-pronounceable consonant string; the last stimulus was always a word. The intervening stimuli were either words or consonant strings depending upon the condition (see Fig. 2). Two types of phrases, in French, were used in the phrase conditions, those containing nouns (NPs) and those containing verbs (VPs). The four-word NPs began with a preposition followed by a possessive pronoun and ended with a noun and adjective, in either order. The four-word VPs began with a proper name followed by an auxiliary verb and ended with a verb and adverb, in either order. We selected eight different word categories in order to maintain an equal amount of category variability both between phrases and between the phrase and list conditions. Also, we chose categories such that all phrase conditions were well formed (i.e. three word possessive-noun-adjective phrases were grammatical constituents, as were noun-adjective pairs). Finally, the categories were chosen to maintain as much parallelism between phrase types as possible, in order to minimize the variability within the full set of phrase conditions, which we averaged together in the analysis. Thus, we equated the general composition functions at each point in both types of phrase – the final two words compose via predicate modification while the initial two words compose via functional application (Kratzer & Heim, 1998).

**Figure 2.**
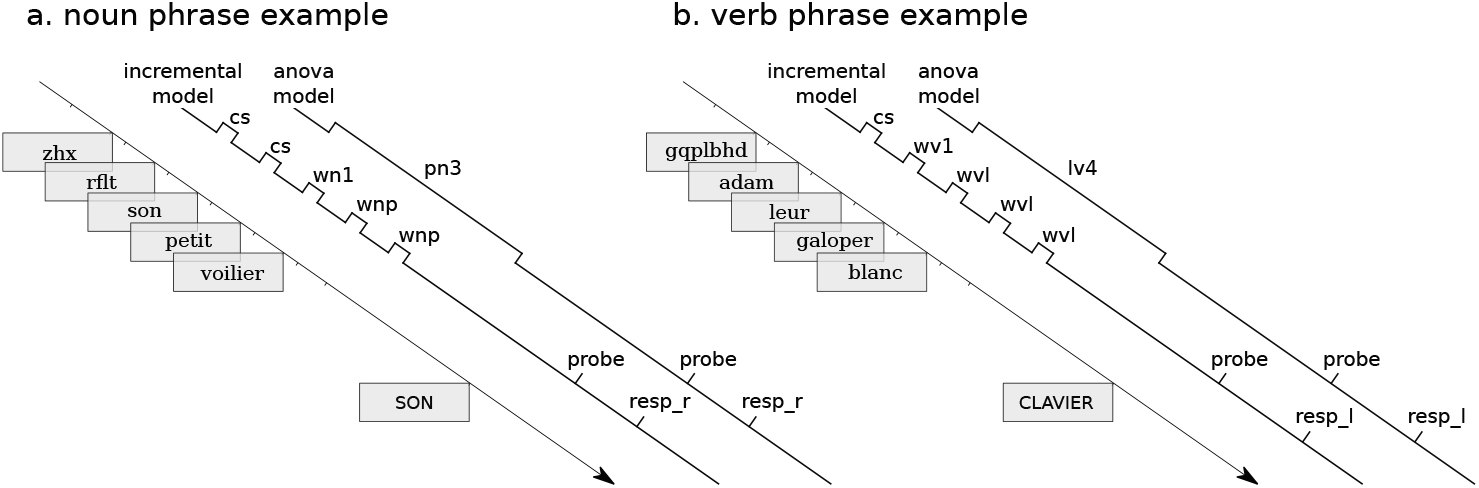
Timelines of two example trials. Each trial consisted in the serial presentation of five items (word or consonant string) at a rate of one every 600ms, followed 2 seconds later by an uppercase item for the probe detection task. Two models (“incremental” and “anova”) were used to analyse fMRI data. The incremental model treated each item as an individual event, from the following types: consonant string (cs), first word in a noun sequence (wn1), word added to a noun phrase (wnp), word added to a noun list (wnl), first word in a verb sequence (wv1), word added to a verb phrase (wvp), word added to a verb list (wvl). The anova model considered each five-item sequence as a single block, of the following types: noun phrases of length 1, 2, 3 and 4 (pn1, pn2, pn3, pn4), verb phrases of length 1, 2, 3 and 4 (pv1, pv2, pv3, pv4), noun lists of length 1, 2, 3 and 4 (ln1, ln2, ln3, ln4), verb lists of length 1, 2, 3 and 4 (lv1, lv2, lv3, lv4). Additionally, both models included events for the display of the probe (probe), and manual responses (right: resp_r, or left: resp_l).

Further, parallel word categories were matched in terms of length, number of syllables, number of phonemes, and (log) frequency. Syllable and phoneme counts were retrieved using http://www.lexique.org/, and frequencies were obtained from the google n-gram corpus (http://storage.googleapis.com/books/ngrams/books/datasetsv2.html). In all cases, we used the least inflected form of each word category, in order to minimize lexical processing. Verbs were always presented in the infinitive form. Only masculine, singular nouns were used, such that the modifying adjective would not need to be inflected. To match orthography between adjectives and adverbs, no adverbs ending in the marked “–ment” suffix were used. To match nouns and verbs, to the greatest degree possible, we used only verbs and nouns ending in “–er”.

Phrases were created by randomly selecting a word from each category (see Supplementary Material: Algorithm to generate the stimuli). The set of words was chosen in order to make any such randomly constructed phrase meaningful. To generate the phrases of length 1, 2 or 3, the length-4 phrases were truncated by replacing the initial words with length-matched consonant strings. Though the selections were random, we required each word to appear equally often in every possible condition. That is, each possessive appeared in an equal number of three and four word conditions; each final adjective appeared in an equal number of one, two, three, and four word conditions. Word lists were constructed by randomly selecting alternating words from categories of different phrase types (see Supplementary Materials). The same requirement regarding an equal number of presentations in each condition for each word was maintained for the lists as well. Finally, probes were selected such that they were present during the preceding words on half of the trials, and on half of the trials they did not appear in the preceding words. For the repeated probes, an equal number were drawn from each possible word position. For example, in the four-word conditions, one-fourth of all correct probes matched the first word. Subjects were instructed to respond with one hand if the probe appeared previously in the trial, and with their other hand if it did not. The hands corresponding to “present” and “not-present” were counter-balanced over subjects, though remained constant for each individual. Trial words were presented in lower-case Georgia font, and sized to subtend between 2° and 4° visual degrees. Probe words were the same size, and presented in upper-case Verdana font. All stimuli were presented using Psychtoolbox (http://psychtoolbox.org/; Brainard, 1997; Pelli, 1997).

All trial types were mixed together and presented in four blocks of 80 trials. Each block contained eight trials of each condition. The conditions were one-, two-, three-, and four-word phrases; one-, two-, three-, and four-word lists; and two additional one-word conditions in which the first word of each two-word condition was displayed in the final position. In the fMRI experiment, there were also the equivalent of two “conditions” of blank trials (i.e. 16 trial length blank screens inserted between the experimental conditions). Trials were presented in pseudo-random order such that all sequential pair orderings were presented equally likely, in order to maximize the power for the fMRI contrasts. Also, sequences were selected such that a given word never appeared in two successive trials.

In addition, 24 twelve-word long sentences and 24 pseudo-words sequences were created to be used in an independent language localizer run.

### Procedure

Before the experiment, subjects completed a practice run outside of the acquisition room. The stimuli for the practice were constructed in order to present each word used in the experiment at least once, in order to minimize repetition effects within the experiment. During the practice run, subjects were warned if they responded too slowly (over two seconds) or incorrectly. They were then told that they would receive no feedback during the experiment.

Each trial began with the presentation of a small (1°) blue cross for 600 ms. This was then followed by the presentation of the five, letter stimuli. Each stimulus was presented for 200 ms and followed by a 400 ms presentation of the cross. After the last word (and subsequent 400 ms break), the cross remained onscreen for 2 seconds, and was followed by the presentation of the probe word (see Fig. 2). The next trial always began 8 seconds following the beginning of the previous trial (giving a maximum of 2400 ms to respond). The probe disappeared when the subject responded, and was removed automatically if the time limit was reached. 20% of the trials were ‘empty’ that is, no stimulus was displayed and participants just had to fixate a cross displayed at the center of the screen. These empty trials were randomly interspersed in the experimental lists to provide a baseline for the models. Subjects were given a self-timed break between each experimental block.

The four experimental runs were followed by a single language localizer run comprising 8 mini-blocks containing 3 sentences and 8 miniblocks containing 3 pseudoword sequences. Mini-blocks lasted 18.4 seconds and were separated by 8 seconds intervals during which a fixation cross was displayed at the center of the screen. Each word or pseudoword was displayed for 300 ms and followed by a blank screen of 100 ms. Sentences or sequences within blocks were separated by 2 seconds. Whenever the word ‘appuyez’ (*press*) or the word ‘cliquez’ (*click*) appeared — events which occurred twice in the run — subjects were instructed to press a response button.

The complete session lasted approximately 1 hour

### MRI data acquisition

For each participant, we first acquired an anatomical scan (T1 weighted image; resolution 1 *×* 1 *×* 1.1mm) and then, for each task block, a series of Echo Planar images with a Time of Repetition of 1.5s and a spatial resolution of 1.5 *×* 1.5 *×* 1.5mm, using a multiband sequence (4 bands) and an iPAT factor set to 2 (Feinberg et al., 2010; Xu et al., 2013).

### MRI data preprocessing

The anatomical images were first defaced with `mask_face` (version 10.15.2018; see https://nrg.wustl.edu/software/face-masking/). Preprocessing was then performed using *fMRIPrep* 1.5.0 (Esteban et al. (2018b); Esteban et al. (2018a); RRID:SCR_016216), which is based on *Nipype* 1.2.2 (Gorgolewski et al. (2011); Gorgolewski et al. (2018); RRID:SCR_002502). Many internal operations of *fMRIPrep* use *Nilearn* 0.5.2 (Abraham et al., 2014a, RRID:SCR_001362), mostly within the functional processing workflow. For more details of the pipeline, see the section corresponding to workflows in *fMRIPrep*’s documentation. The two paragraphs below were copied and pasted from the text automatically generated by fMRIPrep. Although we share in all the derivatives produced by *fMRIPrep* in the public data repository (TODO), only a fraction of those are actually used in our analyses, as detailed in the *Data analysis* section.

#### Anatomical data preprocessing

The T1-weighted (T1w) image was corrected for intensity non-uniformity (INU) with `N4BiasFieldCorrection`(Tustison et al., 2010), distributed with ANTs 2.2.0 (Avants et al., 2008, RRID:SCR_004757), and used as T1w-reference throughout the workflow. The T1w-reference was then skull-stripped with a *Nipype* implementation of the `antsBrainExtraction.sh`workflow (from ANTs), using OASIS30ANTs as target template. Brain tissue segmentation of cerebrospinal fluid (CSF), white-matter (WM) and graymatter (GM) was performed on the brain-extracted T1w using `fast`(FSL 5.0.9, RRID:SCR_002823, Zhang et al., 2001). Volume-based spatial normalization to one standard space (MNI152NLin2009cAsym) was performed through nonlinear registration with `antsRegistration`(ANTs 2.2.0), using brain-extracted versions of both T1w reference and the T1w template. The following template was selected for spatial normalization: *ICBM 152 Nonlinear Asymmetrical template version 2009c* [Fonov et al. (2009), *RRID:SCR_008796; TemplateFlow ID: MNI152NLin2009cAsym]*.

#### Functional data preprocessing

For each of the 5 BOLD runs found per subject (across all tasks and sessions), the following preprocessing was performed. First, a reference volume and its skull-stripped version were generated using a custom methodology of *fMRIPrep*. The BOLD reference was then co-registered to the T1w reference using `flirt`(FSL 5.0.9, Jenkinson & Smith, 2001) with the boundary-based registration (Greve & Fischl, 2009) cost-function. Coregistration was configured with nine degrees of freedom to account for distortions remaining in the BOLD reference. Head-motion parameters with respect to the BOLD reference (transformation matrices, and six corresponding rotation and translation parameters) are estimated before any spatiotemporal filtering using `mcflirt`(FSL 5.0.9, Jenkinson et al., 2002). BOLD runs were slice-time corrected using `3dTshift`from AFNI 20160207 (Cox & Hyde, 1997, RRID:SCR_005927). The BOLD time-series (including slice-timing correction when applied) were resampled onto their original, native space by applying a single, composite transform to correct for head-motion and susceptibility distortions. These resampled BOLD time-series will be referred to as *preprocessed BOLD in original space*, or just *preprocessed BOLD*. The BOLD time-series were resampled into standard space, generating a *preprocessed BOLD run in [‘MNI152NLin2009cAsym’] space*. First, a reference volume and its skull-stripped version were generated using a custom methodology of *fMRIPrep*. Several confounding time-series were calculated based on the *preprocessed BOLD* : framewise displacement (FD), DVARS and three region-wise global signals. FD and DVARS are calculated for each functional run, both using their implementations in *Nipype* (following the definitions by Power et al., 2014). The three global signals are extracted within the CSF, the WM, and the wholebrain masks. Additionally, a set of physiological regressors were extracted to allow for component-based noise correction (*CompCor*, Behzadi et al., 2007). Principal components are estimated after high-pass filtering the *preprocessed BOLD* time-series (using a discrete cosine filter with 128s cut-off) for the two *CompCor* variants: temporal (tCompCor) and anatomical (aCompCor). tCom-pCor components are then calculated from the top 5% variable voxels within a mask covering the subcortical regions. This subcortical mask is obtained by heavily eroding the brain mask, which ensures it does not include cortical GM regions. For aCompCor, components are calculated within the intersection of the aforementioned mask and the union of CSF and WM masks calculated in T1w space, after their projection to the native space of each functional run (using the inverse BOLD-to-T1w transformation). Components are also calculated separately within the WM and CSF masks. For each CompCor decomposition, the *k* components with the largest singular values are retained, such that the retained components’ time series are sufficient to explain 50 percent of variance across the nuisance mask (CSF, WM, combined, or temporal). The remaining components are dropped from consideration. The head-motion estimates calculated in the correction step were also placed within the corresponding confounds file. The confound time series derived from head motion estimates and global signals were expanded with the inclusion of temporal derivatives and quadratic terms for each (Satterthwaite et al., 2013). Frames that exceeded a threshold of 0.5 mm FD or 1.5 standardised DVARS were annotated as motion outliers. All resamplings can be performed with *a single interpolation step* by composing all the pertinent transformations (i.e. head-motion transform matrices, susceptibility distortion correction when available, and co-registrations to anatomical and output spaces). Gridded (volumetric) resamplings were performed using `antsApplyTransforms`(ANTs), configured with Lanczos interpolation to minimize the smoothing effects of other kernels (Lanczos, 1964). Non-gridded (surface) resamplings were performed using `mri_vol2surf`(FreeSurfer).

### Data analysis

All statistical analyses were performed in Python 3 using nilearn (https://nilearn.github.io/) and nistats (https://nistats.github.io/) (Abraham et al., 2014b; Pedregosa et al., 2011).

For the individual subjects analyses, we created a model that associated an event to the presentation of each item (see the “incremental” model in Fig. 2). The types of event were: consonant string (cs), first word in a verb sequence context type (wv1), first word in a noun sequence context type (wn1), added word in either noun list (wnl) or noun phrase (wnp), verb list (wvl) or verb phrase (wvp) context. Three additional variables were added to the design matrix to model events associated to the display of the probe, and the pressing of the left or right response buttons. The 6 rigid motion parameters computed during realignment were also entered in the matrix as variables to control for nuisance effects, as well as the first 6 physiological regressors aCompCor. Functional data were high-pass filtered (cut-off = 128s), detrended using a drift model based on cosine basis functions, and spatially smoothed with a Gaussian kernel with full width at half maximum (fwhm) of 4mm.

At the group level, the analyses consisted of one-sample t-tests performed on individual contrast maps spatially smoothed with a Gaussian filter of fwhm set to 6mm (Mikl et al., 2008).

In addition to the whole-brain analysis, we analyzed the response profiles in seven Regions of Interest selected from previous works on composition: Temporal Pole (TP), anterior Superior Temporal Sulcus (aSTS), posterior Superior Temporal Sulcus (pSTS), Inferior Frontal Gyrus pars orbitalis (IFGorb), Inferior Frontal Gyrus pars triangularis (IFGtri) from Pallier et al. (2011), Inferior Frontal Gyrus pars opercularis (IFGoper) from Zaccarella et al. (2017), and Angular Gyrus/Temporal Parietal Junction (AG/TPJ) from Price et al. (2015). These regions are displayed on Fig. 3. It is important to stress that the names of the ROIs are labels based on localisation of their center’s coordinates on a generic anatomical template and are not meant as precise locations in individual anatomies. For example, given the known inter-individual variability (Amunts et al., 1999), it is inevitable that for some individual, their pars opercularis partially overlap with our IFGtri ROIs.

**Figure 3.**
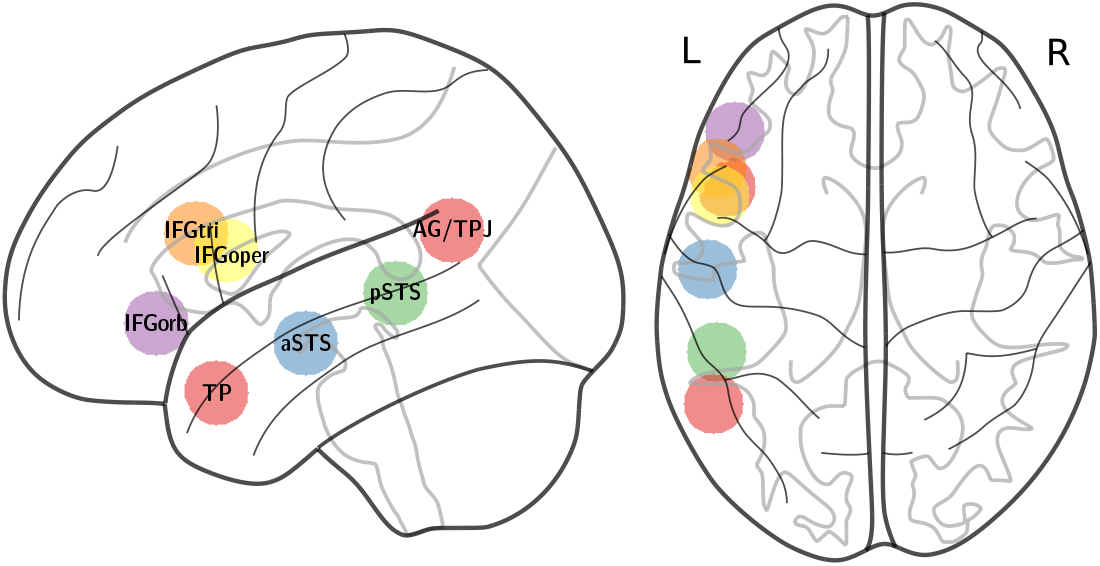
**Regions of Interest** identified in various works on composition. These ROIS were defined as spheres of 10mm radius around the following MNI coordinates: TP (−48, 15, −27), aSTS (−54, −12, −12), pSTS (−51, −39, 3), AG/TPJ (−52, −56, 22), IFGorb (−45, 33, −6), IFGtri (−51, 21, 21), IFGoper (−50, 12, 16)

To sensitize the Region of Interest analyses, we used subject-specific masks within each ROI (Fedorenko et al., 2010). More precisely, for each ROI, we considered for each individual the 300 voxels with the highest z-scores from the contrast between sentences and strings of pseudowords in the localizer run. Because the ROIs contained about 1200 voxels, this represented 25% of the voxels. We used this approach, rather than an absolute threshold on the z-maps, to make sure that each subject contributed equally to the group average activations. Note that the localizer run was independent from the main runs, preventing any problem of circularity.

In order to obtain the detailed profile of response as a function of length, we created an “anova” model featuring 16 variables modeling each of the conditions obtained by crossing the factors Structure (Phrase vs. List), Type (Noun vs Verb) and Length (1 to 4) (see Fig. 2). The stimuli were modeled as boxcars of 2.6 s corresponding to the total display duration of the 5 items sequence. These 16 variables were convolved by the canonical haemodynamic response function of SPM to generate regressors adequate for fMRI analysis. The events associated with probe display and response presses, the 6 motion parameters and the 6 physiological aCompCor regressors, were also included in the design matrix.

## Results

### Analysis of behavioral data

Participants had to perform a probe detection task, that is, to decide after each sequence whether a given word was present in it or not. For each participant and condition, average error rates and reaction times on correct trials were computed, and submitted to repeated measures analyses of variance with the three factors Length (a numerical factor ranging from 1 to 4), Structure (Phrase vs. List) and Type (Noun vs. Verb), using the R package ezANOVA (Lawrence, 2016). These analyses detected a linear effect of Length, reflecting the fact that error rates and reaction times increased with length (see Fig. 4). The effect was marginally significant in the analysis of error rates (*F* (1, 14) = 4.33; *p* = 0.056) and significant in the reaction times analysis (*F* (1, 14) = 65.3; *p* = 1.2 *×* 10^™6^). No other factor nor interaction approaches the 0.05 significance level (see Fig. S1 for detailed results broken down by conditions).

**Figure 4.**
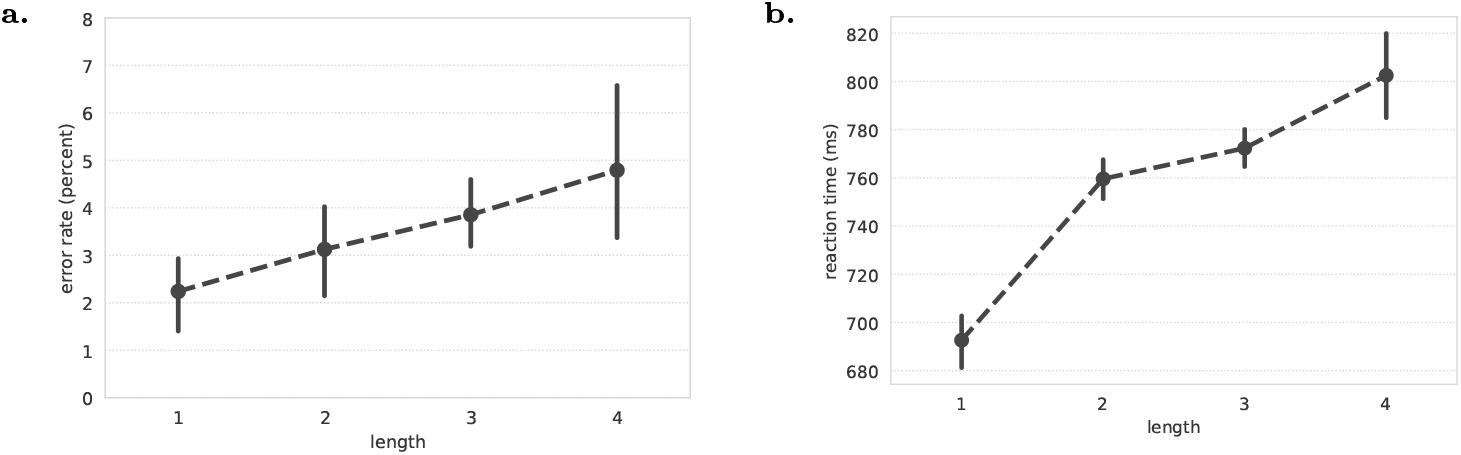
Error rates (a) and reaction times (b) as a function of the Length of the sequence. Bars indicate 95% confidence interval, obtained using bootstrap, after removing the effects of Subject (see Cousineau, 2005).

### Whole-brain analyses

Maps showing the effect of adding a new word to a sequence as a function of Structure (Phrase vs. List) and Type (Nouns vs. Verbs), are displayed on Fig. 5. These maps are presented at uncorrected voxel-wise thresholds *p <* 0.01 and *p <* 0.001 as only few voxels survive a threshold correcting for multiple comparisons (see Supplementary Table S1).

**Figure 5.**
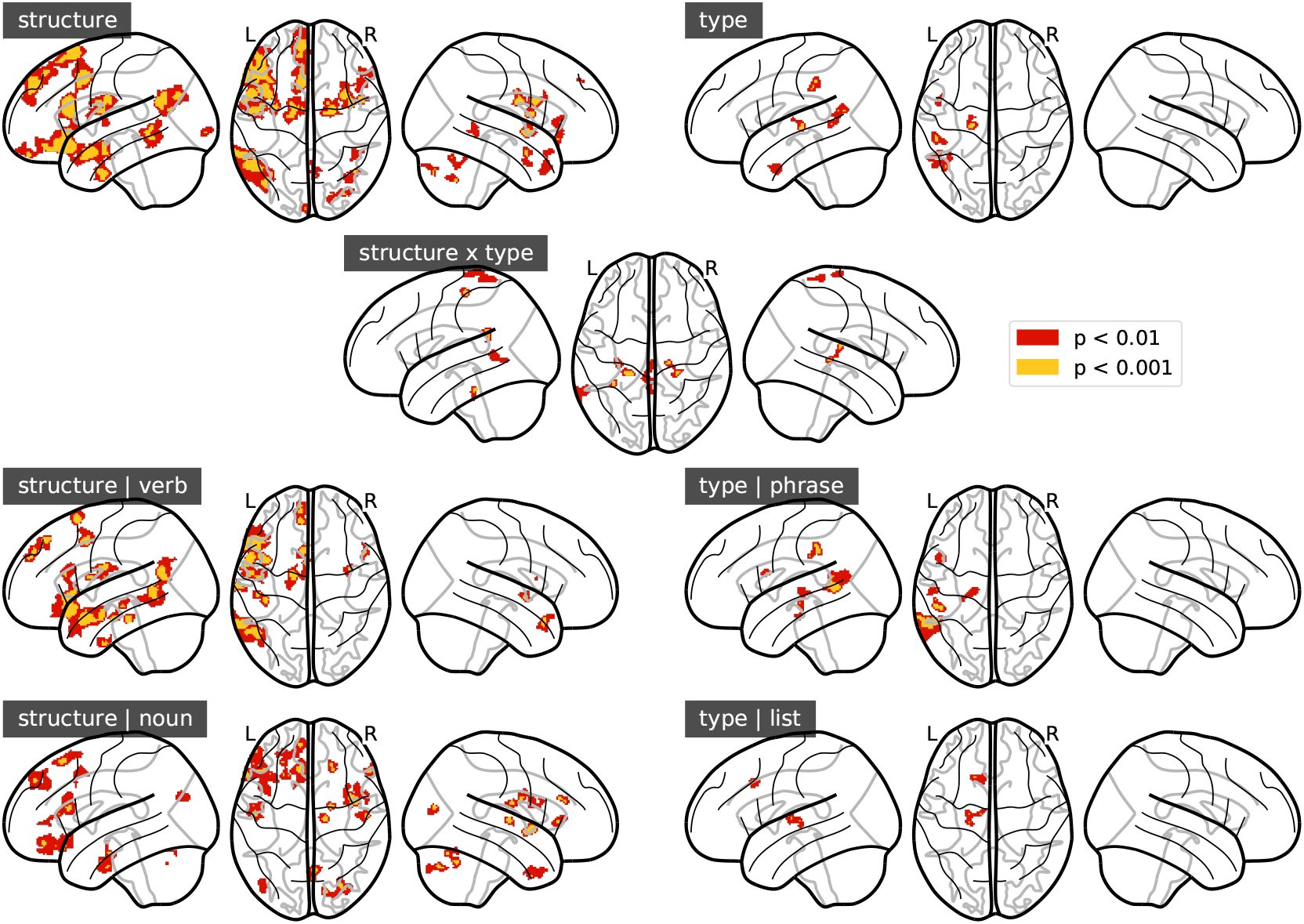
Statistical maps showing activations elicited by the addition of a new word, as a function of Structure (Phrase vs. List) and Type (Verb vs. Noun). Voxels with *p <* 0.01 (uncorrected) and within clusters larger than 100 voxels (voxel size=1.5 *×* 1.5 *×* 1.5mm) are shown in red. Voxels with *p <* 0.001 (uncorrected) are shown in yellow. As few voxels would survive a threshold correcting for multiple comparisons, these maps should be considered as exploratory.

The effect of Structure (top left panel of Fig. 5), contrasting the effects of adding a word to a phrase or to a list, is associated to increased activity in the usual language network comprising regions in the left lateral, and dorsomedial frontal lobes, as well as in the anterior and posterior temporal lobe/angular gyrus.

Further inspection of these maps suggest that verb phrases produce stronger or more robust responses that noun phrases, in posterior temporal regions.

### Regions of Interest

In the temporal and frontal a priori regions of interest displayed on Fig. 3, intersected with the individual language localizer maps, we estimated the effect of adding a word to a sequence as a function of Structure (Phrase vs. List) and Type (Verb vs. Nouns). These data are presented in details in Fig. 6 which shows boxplots of individual effects, along with statistical tests for four relevant contrasts: effect of Structure restricted to Verb or Noun conditions, and effect of Type restricted to Phrases or Lists.

**Figure 6.**
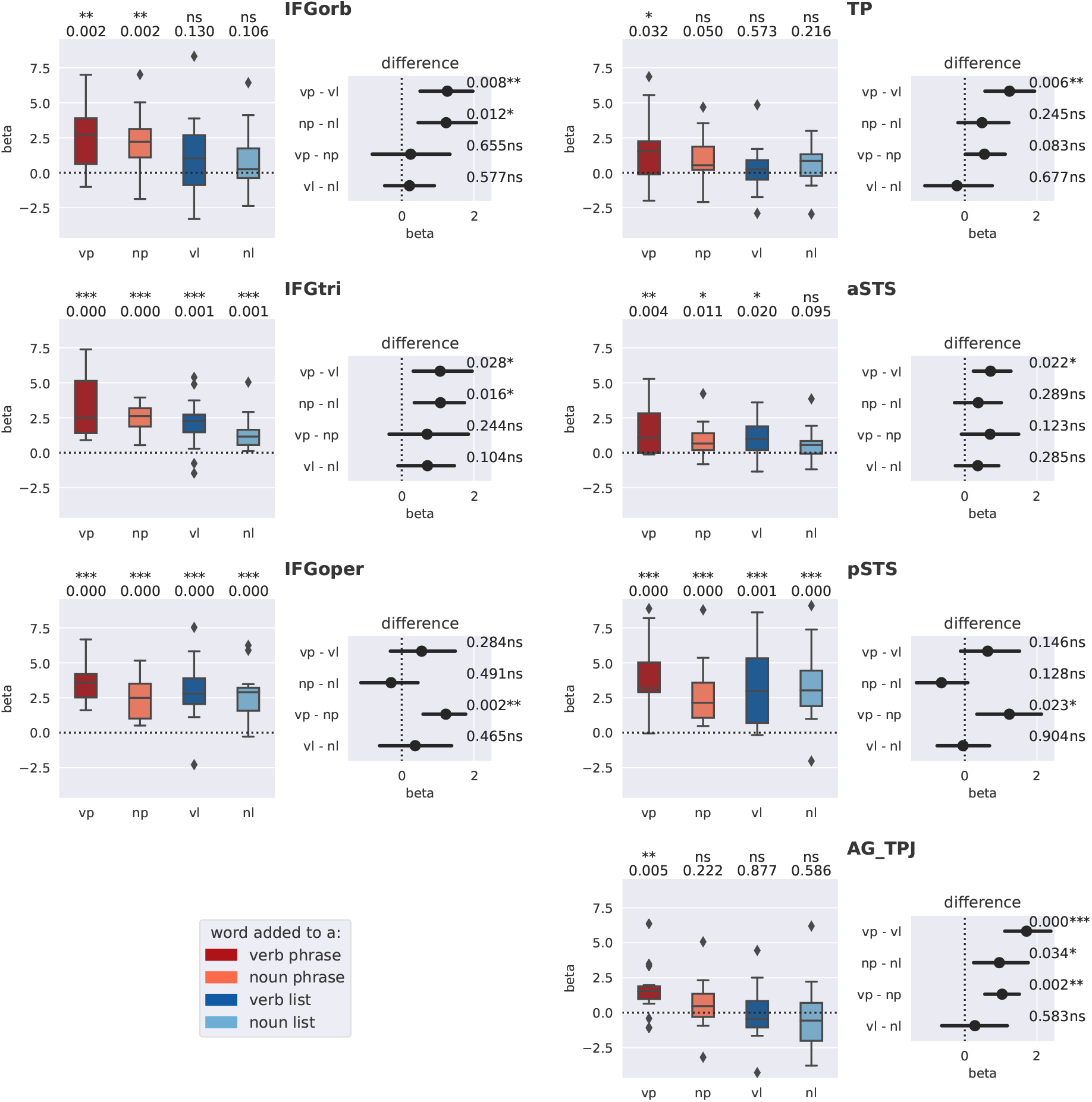
Effect of adding a word to different types of word sequences (vp=verb phrase, np=noun phrase, vl=verb list, nl=noun list), in the a priori regions of interest of Fig. 3. The boxplots show the distributions – across participants – of the effect while the horizontal barplots present the 95% C.I. for contrasts that estimate the effects of Structure for verbs, Structure for nouns, Type for phrases, and Type for lists. Note: p-values are not corrected for multiple comparisons. As there are 8 ∗ 7 = 57 tests on this page, the Bonferroni 5% threshold is *p* ≃ 0.001, so only the tests with ‘***’ would pass it.

All of the ROIs show an increase in activation when a word is added to a verb phrase (dark red boxplots), and, in the case of noun phrases (light red boxplots), adding a word produces an increase everywhere except in the AG/TPJ. These plots also reveal that the effect of adding an item to a word sequence is significantly stronger in verb phrases than in noun phrases in IFGoper, pSTS and AG/TPJ.

A subset of regions also show significant increases when a word is added to a list (blue boxplots): these are IFGoper, IFGtri and pSTS. In IFGoper and in pSTS regions, adding a word to a phrase or to a list produces an increase of similar amplitude, while in IFGtri, the effect is stronger in phrases than in lists.

Fig. 7 shows the profile of response to Length (number of words in each sequence, ranging from 1 to 4), estimated from the “anova” model. The results of the statistical tests are in general agreement with Fig.6. Nevertheless, one can see that the effect of Length is not perfectly linear. In AG/TPJ, aSTS, IFGorb and IFGtri, activation receives a boost in the condition Verb phrase - Length 4.

**Figure 7.**
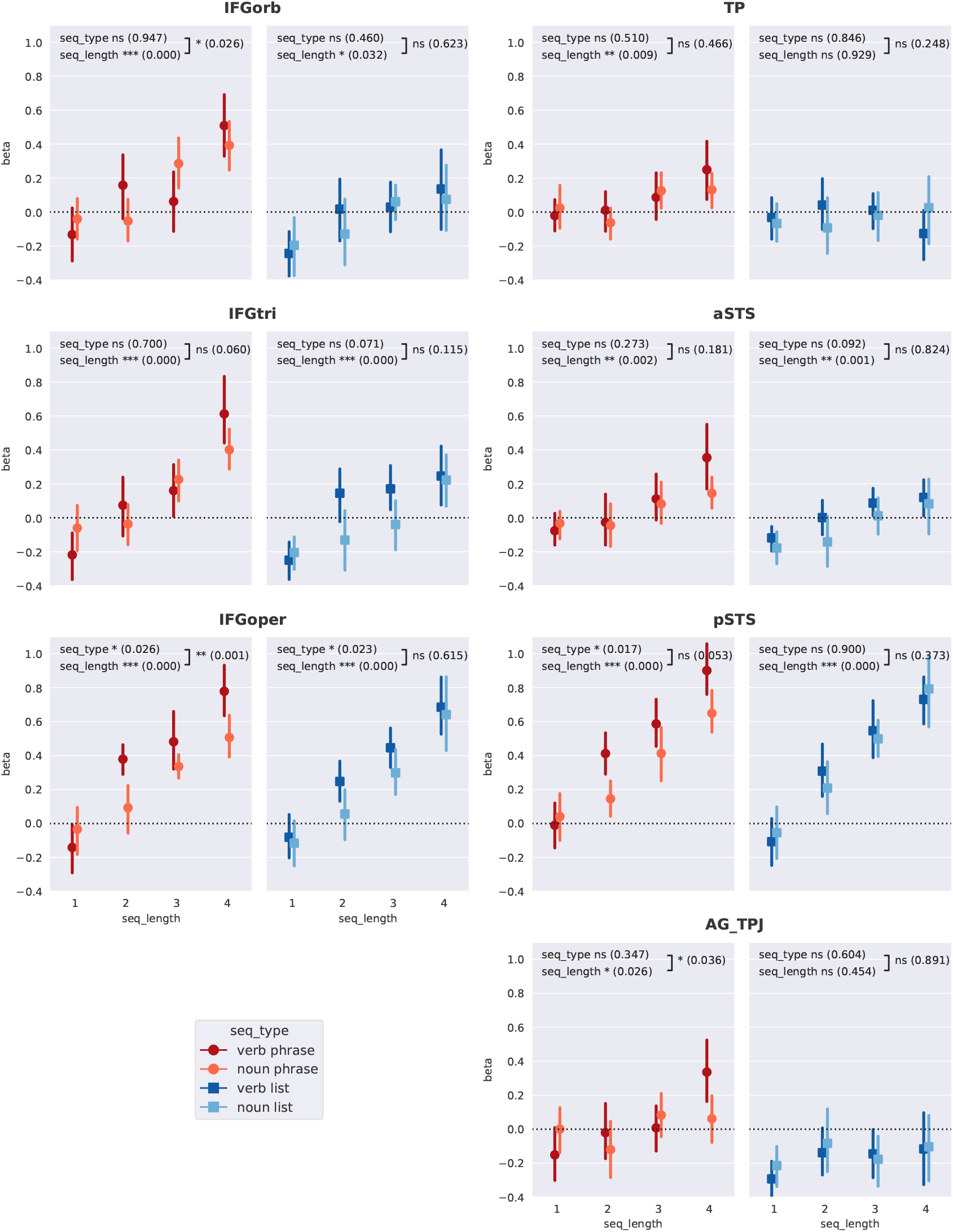
Effect of Length in Regions of Interest. Responses estimated from the Anova model crossing Length, Structure and Type. Error bars represent 95% confidence intervals computed by bootstrap after removing the m1a5in effects of Subjects (Cousineau, 2005). The p-values test for the effect of Type, Length and their interaction (they are not corrected for the 3*2*7= 42 tests displayed on the figure).

### Interactions between regions and conditions

To assess if the response patterns differed between regions, we computed the interactions between Regions (grouped by pairs), and the four contrasts reported on Fig. 6: the effects of Structure, restricted to verbs and to nouns, and the effects of Type restricted to phrases and to lists. Table 1 reports those tests. This analysis reveals that the pSTS (last column) has a pattern of responses that differ significantly from the other regions except IFGoper; IFGoper also stands out, differing from all regions except pSTS and aSTS. This is essentially due to the strong responses to Length in Lists for these two regions, IFGoper and pSTS (see Fig. 7).

**Table 1:** Interaction tests between pairs of Regions of Interest and the contrasts on Structure and Type. For example, the top left p-value shows that regions IFGoper and AG/TPJ have a significant (p=.023) difference in the effect of Structure (phrase vs list) for verbs. Note: these interactions were computed from the incremental model. Note that the p-values are not corrected for the 21 *4 = 84 tests presented here; no test survives a Bonferroni correction (alpha=.05/84=0.0006).

|  | IFGoper | IFGorb | IFGtri | TP | aSTS | pSTS |  |
| --- | --- | --- | --- | --- | --- | --- | --- |
| AG/TPJ | 0.023 * | 0.201 ns | 0.131 ns | 0.354 ns | 0.045 * | 0.061 ns | structure / verbs |
|  | 0.004 ** | 0.615 ns | 0.791 ns | 0.085 ns | 0.065 ns | 0.001 *** | structure / nouns |
|  | 0.588 ns | 0.128 ns | 0.562 ns | 0.165 ns | 0.404 ns | 0.698 ns | type / phrases |
|  | 0.815 ns | 0.837 ns | 0.220 ns | 0.171 ns | 0.866 ns | 0.476 ns | type / lists |
| IFGoper |  | 0.098 ns | 0.172 ns | 0.131 ns | 0.742 ns | 0.868 ns | structure / verbs |
|  |  | 0.004 ** | 0.001 ** | 0.092 ns | 0.083 ns | 0.384 ns | structure / nouns |
|  |  | 0.074 ns | 0.252 ns | 0.009 ** | 0.125 ns | 0.977 ns | type / phrases |
|  |  | 0.672 ns | 0.071 ns | 0.290 ns | 0.979 ns | 0.447 ns | type / lists |
| IFGorb |  |  | 0.662 ns | 0.946 ns | 0.234 ns | 0.258 ns | structure / verbs |
|  |  |  | 0.709 ns | 0.170 ns | 0.071 ns | 0.001 *** | structure / nouns |
|  |  |  | 0.365 ns | 0.523 ns | 0.214 ns | 0.101 ns | type / phrases |
|  |  |  | 0.082 ns | 0.232 ns | 0.681 ns | 0.428 ns | type / lists |
| IFGtri |  |  |  | 0.718 ns | 0.471 ns | 0.394 ns | structure / verbs |
|  |  |  |  | 0.252 ns | 0.085 ns | 0.000 *** | structure / nouns |
|  |  |  |  | 0.711 ns | 0.993 ns | 0.291 ns | type / phrases |
|  |  |  |  | 0.088 ns | 0.472 ns | 0.135 ns | type / lists |
| TP |  |  |  |  | 0.132 ns | 0.246 ns | structure / verbs |
|  |  |  |  |  | 0.706 ns | 0.017 * | structure / nouns |
|  |  |  |  |  | 0.488 ns | 0.053 ns | type / phrases |
|  |  |  |  |  | 0.143 ns | 0.677 ns | type / lists |
| aSTS |  |  |  |  |  | 0.865 ns | structure / verbs |
|  |  |  |  |  |  | 0.002 ** | structure / nouns |
|  |  |  |  |  |  | 0.177 ns | type / phrases |
|  |  |  |  |  |  | 0.164 ns | type / lists |

A limit of this type of analysis should be stressed: because different regions can have different global strength of responses, an interaction could simply be due to the fact that one region responds more strongly than the other. It is therefore important to look at Fig. 6 and 7 to interpret these tests.

## Discussion

The present fMRI experiment was designed as a follow-up of Pallier et al. (2011) and Bemis & Pylkkänen (2013) using more tightly controlled stimuli. Namely, we investigated combinatorial activity by presenting short visual sequences (1 to 4 words long) forming either meaningful grammatical phrases (e.g. “avec son beau clavier”, *with his nice keyboard*) or unstructured lists (“Adam son plus clavier”, *Adam his more keyboard*). As in Bemis & Pylkkänen (2013), the sequences were padded with consonant strings in order to equalize their length, and to avoid the juxtaposition of several short constituents that occurred in the materials of Pallier et al. (2011). Moreover, the procedure to generate unstructured lists was such that neighboring words were less likely to merge here than in the previous studies. In the current experiment, the list stimuli were constructed by mixing words coming from syntactic categories that cannot be recombined even after “mentally scrambling” the words (for example, “sur doit boîtier tôt”: *above must box early*). This blocks potential attempts to “repair” the sentence by scrambling (Mollica et al., 2018).

We also better controlled the phrase condition: half of the stimuli were built around nouns, while the other half were built around verbs (see Fig. 1 and Supplementary Materials for the full set of stimuli). Compared to the study by Pallier et al., there was a probe monitoring task after each sequence to ensure that participants paid attention until the end of every sequence.

The main findings of the current study can be summarized as follow: (a) adding a word to a growing phrase structure yields an additional activation in multiple areas of the language network, comprising the perisylvian areas (IFG, temporal regions and angular gyrus), and more dorsal regions (dorsomedial prefrontal cortex, (pre)-SMA, premotor cortex); (b) some regions, pSTS, IFGoper and, to a lesser extent IFGtri, also increase in activity when a word is added to a unstructured list; (c) adding words to verb phrases typically yields stronger activations than adding words to noun phrases.

These results are consistent with Pallier et al. (2011). The only qualitative difference is the fact that the pSTS, which showed a significant effect of Structure in Pallier et al., did not show such an effect in the current study: it responded as strongly to lists of words than to well-formed phrases. Yet, the present study showed an effect of Structure when the analyses were restricted to the verb condition. As in Pallier et al, the condition with the longest constituent (c12) was always a complete sentence containing a finite verb, the two studies are consistent. It is noteworthy that the pSTS and the IFGoper strongly responded to unstructured word lists, in both studies, and even in the Jabberwocky condition in Pallier et al (2011), a fact that we will discuss below.

The current experiment did not include a Jabberwocky condition. In Pallier et al. (2011) and Matchin et al. (2017), IFGorb behaved differently from the anterior temporal regions. To further characterize the behavior of these regions, we reviewed a series of studies which compared responses to sentences, or to Jabber-worky, to word or non-word lists Price et al. (2015); Bemis & Pylkkänen (2013); Humphries et al. (2006); Matchin et al. (2017); Vandenberghe et al. (2002). The consistent observations can be summarized as follows: (a) TP, AG/TPJ and aSTS are essentially sensitive to semantic manipulations: they systematically show stronger activation in sentences than in scrambled sentences. (b) IFGtri, IFGoper and pSTS respond to any linguistic materials, whether it is structured or not, whether they contain real or pseudowords. Yet, they typically respond more strongly if the content is structured; (c) IFGorb has a weaker response to unstructured stimuli than the other parts of the IFG and the pSTS. Moreover, contrary to TP, aSTS and AG/TPJ, IFGorb is sensitive to structure even with Jabberwocky material.

### The roles of IFGoper, pSTS and IFGtri

Matchin et al. (2017) proposed that IFGoper is involved in Working Memory (the rehearsal network). Our observations are compatible with this notion. Indeed, we used a probe task which required trying to remember the 1-4 words in both lists and phrases. Working memory areas are expected to increase their responses with the number of stored items (Narayanan et al., 2005; D’Esposito & Postle, 2015)), regardless of condition, exactly as we found.

Matchin even observed stronger activation for lists than for phrases or sentences. Even though we used the same probe detection task as Matchin et al., we did not observe a stronger length effect for lists than for sentences in any of the regions of interest. In the case of the IFGoper, we found a trend towards stronger activation for verb phrases than verb lists. This does not preclude the possibility that IFGoper participates in the rehearsal loop, but demonstrates that it may be sensitive to other factors than just memory load. As noted in the introduction, the IFG and the pSTS have been detected in many studies of syntactic processing (Rodd et al., 2010, e.g. when manipulating syntactic ambiguity).

On the other hand, Goucha & Friederici (2015) and Zaccarella et al. (2017) claimed that IFGoper or, rather, a subpart of it, is performing the pure syntactic ‘merge’ operation, while IFGtri and pSTS would be involved when derivational morphology processes are implicated. Most of the nouns and verbs that we used were multimorphemic — which is unavoidable for verb in French.

The literature does rarely show clear-cut, systematic, functional dissociations between IFGoper, pSTS and IFGtri. The IFGtri always shows an effect of sentences vs. words lists, while the effect is more variable in IFGoper and pSTS, as these regions typically respond more strongly to word lists (relative to baseline) than the IFGtri. Nevertheless, this does not preclude the possibility that these regions are involved in storing items in verbal working memory in distinct formats, for instance a phonological loop for IFGoper versus a lexico-syntactic loop for IFGtri.

Matchin et al. (2017) argued that IFGtri and pSTS are involved in syntactic prediction, based on the fact that they observed significant effects for sentences vs. word lists, but no effect for simple phrases vs. word lists. Here, we observed that these regions — IFGtri/oper and pSTS — responded strongly to unstructured lists of words. This could be partly due to lexical processes (e.g. word recognition). However, their responses are not purely driven by the number of words : verb phrases yield stronger activation than noun phrases. As we have matched the verbs and the nouns on form properties (length in characters and lexical frequency), this difference is likely due to difference in the syntactic or semantic properties of the verbs and nouns.

At this stage, it must be stressed that stronger responses to verbs than to nouns have been repeatedly reported in the IFG, pSTS and AG/TPJ regions in the literature. Tyler et al. (2008) reported stronger responses in the pSTS/pMTG to verb phrases vs. noun phrases (“they visit” vs. “the visit”), corresponding roughly to our contrast Type / Phrase. Longe et al. (2007) found that inflecting verbs yields more activation than inflecting nouns: they reported an interaction “snails, hear*s*” vs. “snail, hear” in the pSTS/MTG. Similarily, Willms et al. (2011) found stronger activations in pSTS and frontal areas when producing inflected verbs rather than inflected nouns. Palti et al. (2007) reported stronger activation in the pSTS and pIFG for verbs than for nouns.

One potential explanation for the stronger responses to verbs is that verbs typically project more structure than nouns, for example in terms of argument structure. Consistent with this interpretation, Hernández et al. (2014) found that transitive verbs provoked stronger activations than intransitive ones in the pSTS and IFG. In the same vein, Thompson et al. (2007) have reported more activation for two- and three-arguments verbs compared to one-argument verbs in the AG/TPJ.

In our study, it is also noteworthy that the longer phrases in the verb condition were actually complete sentences (“Marc doit rouler vite”, *Marc must drive fast*). We observed an especially strong increase from length 3 to length 4, only for the verb condition, in the three regions of the IFG and in the aSTS. The difference between nouns and verbs in our study can thus be partly driven by a difference between full sentences, and noun phrases. Note that in some formalisms, e.g. generative grammars, a sentence require more nodes (e.g. a tense node) than a simple noun phrase. From this point of view, the activation might reflect the linguistic complexity of the object represented, as measured by the number of tree nodes that it entails.

Pallier et al. discovered that adding a word to a growing phrase structure imposes an additional activation in multiple areas of the language network (see also Nelson et al., 2017; Fedorenko et al., 2016). The present study strengthens the previous findings by using a tighter design, making sure that subjects paid attention to the entire sequence, and focusing on short constituents (size 2 to 4). Together with results from other publications, it shows that even if most of the language network is sensitive to structure, different sub regions have different response profiles to lists of words or sentences. Trying to understand the precise computations performed by those regions remains a valuable goal that many teams are actively pursuing.

## Data and code availability

The raw dataset is available at https://openneuro.org/datasets/ds002905. The Python 3 source code for the analysis is available on GitHub: https://github.com/l-bg/simple_composition_fmri.

## Supporting information

Supplementary Material: Algorithm to generate the stimuli.

Supplementary Figure S1: Error rates (a) and reaction times (b), broken down by conditions (S: Structure, T: Type, L: Length) and corresponding ANOVA tables.

Supplementary Table S1: Results of whole-brain analyses from the incremental model.

## CRediT author statement

Laurent Bonnasse-Gahot: Methodology, Software, Formal Analysis, Visualization, Writing; Doug Bemis: Conceptualization, Investigation, Software; Martin Perez-Guevara: Data Curation; Stanislas Dehaene: Conceptualization, Writing Review & Editing; Christophe Pallier: Conceptualization, Methodology, Software, Writing, Project administration, Funding acquisition.

## Acknowledgments

We thank Chantal Ginisty, Séverine Desmidt, Séverine Roger and Adrien Desbois for help in data acquisition, and Antoine Grigis for the defacing of anatomical images. This research was supported by the French National Research Agency (ANR Programme blanc, project “Construct”, 140301).

