## Supplementary Materials for "An fMRI study of composition in noun and verb phrases"

### Supplementary Material: Algorithm to generate the stimuli

The stimuli were sequences obtained by combining French words (see English glosses infra) in the following ways:

#### Phrase condition

##### Noun phrase condition

$$\left\{\begin{array}{c}\text{avec}\\\text{dans}\\\text{pour}\\\text{sans}\\\text{sous}\\\text{sur}\\\text{vers}\end{array}\right\}_{\text{P}} \times \left\{\begin{array}{c}\text{leur}\\\text{mon}\\\text{son}\end{array}\right\}_{\text{DET}} \times \left\{\begin{array}{c}\text{beau}\\\text{grand}\\\text{nouveau}\\\text{petit}\\\text{vieux}\end{array}\right\}_{\text{ADJ}} \times \left\{\begin{array}{c}\text{boîtier}\\\text{casier}\\\text{cendrier}\\\text{clavier}\\\text{grenier}\\\text{panier}\\\text{sablier}\\\text{rocher}\\\text{tablier}\\\text{voilier}\end{array}\right\}_{\text{N}}$$

$$\left\{\begin{array}{c}\text{avec}\\\text{dans}\\\text{pour}\\\text{sans}\\\text{sous}\\\text{sur}\\\text{vers}\end{array}\right\}_{\text{P}} \times \left\{\begin{array}{c}\text{leur}\\\text{mon}\\\text{son}\end{array}\right\}_{\text{DET}} \times \left\{\begin{array}{c}\text{boîtier}\\\text{casier}\\\text{cendrier}\\\text{clavier}\\\text{grenier}\\\text{panier}\\\text{sablier}\\\text{rocher}\\\text{tablier}\\\text{voilier}\end{array}\right\}_{\text{N}} \times \left\{\begin{array}{c}\text{blanc}\\\text{bleu}\\\text{noir}\\\text{rouge}\\\text{vert}\end{array}\right\}_{\text{ADJ}}$$

##### Verb phrase condition

$$\left\{\begin{array}{c}\text{Adam}\\\text{Paul}\\\text{Yves}\\\text{Marc}\\\text{Jean}\\\text{Luc}\\\text{Jade}\end{array}\right\}_{\text{PN}} \times \left\{\begin{array}{c}\text{doit}\\\text{peut}\\\text{va}\end{array}\right\}_{\text{MOD}} \times \left\{\begin{array}{c}\text{mieux}\\\text{trop}\\\text{beaucoup}\\\text{plus}\\\text{moins}\end{array}\right\}_{\text{ADV}} \times \left\{\begin{array}{c}\text{jongler}\\\text{nager}\\\text{patiner}\\\text{plonger}\\\text{chasser}\\\text{rouler}\\\text{parader}\\\text{danser}\\\text{voyager}\\\text{galoper}\end{array}\right\}_{\text{V}}$$

$$\left\{\begin{array}{c}\text{Adam}\\\text{Paul}\\\text{Yves}\\\text{Marc}\\\text{Jean}\\\text{panier Luc}\\\text{Jade}\end{array}\right\}_{\text{PN}} \times \left\{\begin{array}{c}\text{doit}\\\text{peut}\\\text{va}\end{array}\right\}_{\text{MOD}} \times \left\{\begin{array}{c}\text{jongler}\\\text{nager}\\\text{patiner}\\\text{plonger}\\\text{chasser}\\\text{rouler}\\\text{parader}\\\text{danser}\\\text{voyager}\\\text{galoper}\end{array}\right\}_{\text{V}} \times \left\{\begin{array}{c}\text{vite}\\\text{tôt}\\\text{tard}\\\text{ainsi}\\\text{loin}\end{array}\right\}_{\text{ADV}}$$

### List condition

#### Noun list condition

$$\left\{\begin{array}{c}\text{Adam}\\\text{Paul}\\\text{Yves}\\\text{Marc}\\\text{Jean}\\\text{Luc}\\\text{Jade}\end{array}\right\}_{\text{PN}} \times \left\{\begin{array}{c}\text{leur}\\\text{mon}\\\text{son}\end{array}\right\}_{\text{DET}} \times \left\{\begin{array}{c}\text{mieux}\\\text{trop}\\\text{beaucoup}\\\text{plus}\\\text{moins}\end{array}\right\}_{\text{ADV}} \times \left\{\begin{array}{c}\text{boîtier}\\\text{casier}\\\text{cendrier}\\\text{clavier}\\\text{grenier}\\\text{panier}\\\text{sablier}\\\text{rocher}\\\text{tablier}\\\text{voilier}\end{array}\right\}_{\text{N}}$$

$$\left\{\begin{array}{c}\text{avec}\\\text{dans}\\\text{pour}\\\text{sans}\\\text{sous}\\\text{sur}\\\text{vers}\end{array}\right\}_{\text{P}} \times \left\{\begin{array}{c}\text{doit}\\\text{peut}\\\text{va}\end{array}\right\}_{\text{MOD}} \times \left\{\begin{array}{c}\text{boîtier}\\\text{casier}\\\text{cendrier}\\\text{clavier}\\\text{grenier}\\\text{panier}\\\text{sablier}\\\text{rocher}\\\text{tablier}\\\text{voilier}\end{array}\right\}_{\text{N}} \times \left\{\begin{array}{c}\text{vite}\\\text{tôt}\\\text{tard}\\\text{ainsi}\\\text{loin}\end{array}\right\}_{\text{ADV}}$$

#### Verb list condition

$$\left\{\begin{array}{c}\text{Adam}\\\text{Paul}\\\text{Yves}\\\text{Marc}\\\text{Jean}\\\text{Luc}\\\text{Jade}\end{array}\right\}_{\text{PN}} \times \left\{\begin{array}{c}\text{leur}\\\text{mon}\\\text{son}\end{array}\right\}_{\text{DET}} \times \left\{\begin{array}{c}\text{jongler}\\\text{nager}\\\text{patiner}\\\text{plonger}\\\text{chasser}\\\text{rouler}\\\text{parader}\\\text{danser}\\\text{voyager}\\\text{galoper}\end{array}\right\}_{\text{V}} \times \left\{\begin{array}{c}\text{blanc}\\\text{bleu}\\\text{noir}\\\text{rouge}\\\text{vert}\end{array}\right\}_{\text{ADJ}}$$

$$\left\{\begin{array}{c}\text{avec}\\\text{dans}\\\text{pour}\\\text{sans}\\\text{sous}\\\text{sur}\\\text{vers}\end{array}\right\}_{\text{P}} \times \left\{\begin{array}{c}\text{doit}\\\text{peut}\\\text{va}\end{array}\right\}_{\text{MOD}} \times \left\{\begin{array}{c}\text{beau}\\\text{grand}\\\text{nouveau}\\\text{petit}\\\text{vieux}\end{array}\right\}_{\text{ADJ}} \times \left\{\begin{array}{c}\text{jongler}\\\text{nager}\\\text{patiner}\\\text{plonger}\\\text{chasser}\\\text{rouler}\\\text{parader}\\\text{danser}\\\text{voyager}\\\text{galoper}\end{array}\right\}_{\text{V}}$$

#### English glosses

**Prepositions (P):** avec *with*, dans *in*, pour *for*, sans *without*, sous *under*, sur *on*, vers *towards*

**Determiners (DET):** leur *their*, mon *my*, son *his*

**Adjectives (ADJ):** beau *nice*, grand *big*, nouveau *new*, petit *small*, vieux *old*, blanc *white*, bleu *blue*, noir *black*, rouge *red*, vert *green*

**Nouns (N):** boîtier *box*, casier *case*, cendrier *hashtray*, clavier *keyboard*, grenier *attic*, panier *basket*, sablier *hourglass*, rocher *rock*, tablier *apron*, voilier *sail-ship*

**Proper names (PN):** Adam, Paul, Yves, Marc, Jean, Luc, Jade

**Modifiers/Auxiliaries (MOD):** doit *must*, peut *can*, va *is going to*

**Adverbs (ADV):** mieux *better*, trop *too much*, beaucoup *a lot*, plus *more*, moins *less*, vite *quickly*, tôt *early*, tard *late*, ainsi *thus*, loin *far*,

**Verbs (V):** jongler *to juggle*, nager *to swim*, patiner *to skate*, plonger *to dive*, chasser *to hunt*, rouler *to drive*, parader *to parade*, danser *to dance*, voyager *to travel*, galoper *to gallop*

**a.**

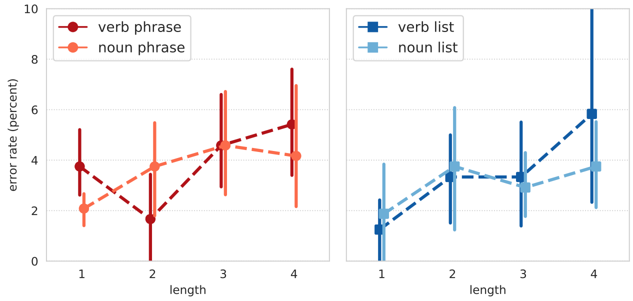


| effect | F(1,14) | $p$ | $\eta^2_G$ |
|---|---|---|---|
| S | 0.46 | 0.51 | 6.59e-3 |
| T | 0.28 | 0.61 | 2.22e-3 |
| L | 4.33 | 0.06 | 8.69e-2 |
| S×T | 0.03 | 0.87 | 1.65e-4 |
| S×L | 0.07 | 0.80 | 1.06e-3 |
| T×L | 2.42 | 1.42 | 8.05e-3 |
| S×T×L | 3.16 | 0.10 | 5.56e-3 |

**b.**

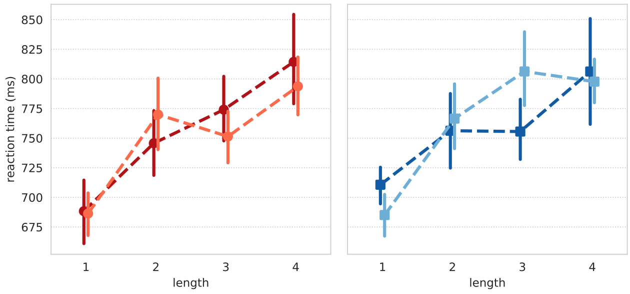


| effect | F(1,14) | $p$ | $\eta^2_G$ |
|---|---|---|---|
| S | 1.42 | 0.25 | 8.67e-3 |
| T | 0.004 | 0.95 | 8.08e-5 |
| L | 65.3 | 1.2e-6* | 4.75e-1 |
| S×T | 0.67 | 0.43 | 5.46-3 |
| S×L | 0.10 | 0.76 | 1.03e-3 |
| T×L | 0.001 | 0.91 | 6.1e-5 |
| S×T×L | 2.37 | 0.15 | 1.78e-2 |

Figure S1: **Error rates (a) and reaction times (b), broken down by conditions (S: Structure, T: Type, L: Length) and corresponding ANOVA tables.** Bars indicate 95% confidence interval, obtained using bootstrap, after removing the effects of Subject (see Cousineau, 2005). The column $\eta^2_G$ provides the generalized eta-squared effect size (Olejnik and Algina, 2003).

| contrast | region | BA | x | y | z | z-value | cluster size |
|---|---|---|---|---|---|---|---|
| *structure* | preSMA | 6 | -7 | 18 | 64 | 4.18 | 938 |
| | dmPFC | 8 | -7 | 27 | 60 | 3.30 | |
| | dmPFC | 9 | -7 | 52 | 37 | 4.72 | 897 |
| | LPMC | 6 | -43 | 10 | 43 | 4.31 | 1036 |
| | LPMC | 6 | -39 | 13 | 37 | 3.86 | |
| | IFGtri | 45 | -60 | 21 | 15 | 4.45 | 1869 |
| | IFGtri | 45 | -43 | 24 | 13 | 3.74 | |
| | IFGoper | 44 | -36 | 19 | 19 | 3.52 | |
| | dlPFC | 46 | -57 | 25 | 22 | 3.68 | |
| | IFGorb | 47 | -45 | 27 | -18 | 4.60 | 1869 |
| | IFGorb | 47 | -43 | 36 | -19 | 4.30 | |
| | IFGorb | 47 | -49 | 19 | -16 | 3.94 | |
| | IFGorb | 47 | -51 | 24 | -7 | 3.93 | |
| | LCaudate | 48 | -16 | -10 | 25 | 4.10 | 384 |
| | LCaudate | 48 | -16 | -3 | 21 | 3.49 | |
| | TP | 38 | -60 | 3 | -21 | 4.60 | 756 |
| | TP | 38 | -51 | 7 | -24 | 3.80 | |
| | AG/TPJ | 39 | -54 | -55 | 21 | 4.78 | 1154 |
| | AG/TPJ | 39 | -43 | -58 | 18 | 3.23 | |
| *structure* \| *verb* | LPMC | 6 | -37 | 6 | 45 | 4.11 | 351 |
| | LPMC | 6 | -45 | 10 | 43 | 3.66 | |
| | IFGorb | 47 | -45 | 25 | -18 | 3.99 | 1289 |
| | IFGtri | 45 | -54 | 21 | -4 | 3.95 | |
| | IFGorb | 47 | -48 | 25 | -10 | 3.61 | |
| | TP | 38 | -57 | 9 | -18 | 4.64 | 722 |
| | pSTS | 21 | -63 | -49 | 3 | 4.32 | 742 |
| | AG/TPJ | 39 | -46 | -61 | 13 | 3.87 | 654 |
| | AG/TPJ | 39 | -49 | -55 | 19 | 3.64 | |
| *type* \| *phrase* | pSTS | 21 | -57 | -51 | 4 | 4.16 | 631 |
| | pSTS | 21 | -66 | -43 | 6 | 3.22 | |

Table S1: **Results of whole-brain analyses from the incremental model**. The maps were thresholded at $p < 0.001$ voxel-wise, with a cluster threshold of 100 voxels (uncorrected; the FWE-corrected threshold and FDR-corrected threshold for cluster size are 492 voxels, computed using SPM). Only coordinates of peaks that are at least 8mm apart are reported. Cluster size are expressed in $mm^3$. The Brodmann labels (BA) were estimated using the "mni2tal" tool at `http://sprout022.sprout.yale.edu/mni2tal/mni2tal.html`. LPMC=Left Premotor Cortex; dmPFC: dorso-medial Prefrontal Cortex; dlPFC: dorso-lateral Prefrontal Cortex; preSMA=pre-Supplementary Motor Area.
